# Analysis of single nucleotide polymorphisms between 2019-nCoV genomes and its impact on codon usage

**DOI:** 10.1101/2020.08.05.237404

**Authors:** Suruchi Gupta, Ravail Singh, Prosenjit Paul

## Abstract

The spread of COVID-19 is a global concern that has taken a toll on entire human health. Researchers across the globe has been working in devising the strategies to combat this dreadful disease. Studies focused on genetic variability helps design effective drugs and vaccines. Considering this, the present study entails the information regarding the genome-wide mutations detected in the 108 SARS CoV-2 genomes worldwide. We identified a few hypervariable regions localized in orf1ab, spike, and nucleocapsid gene. These nucleotide polymorphisms demonstrated their effect on both codon usage as well as amino acid usage pattern. Altogether the present study provides valuable information that would be helpful to ongoing research on 2019-nCoV vaccine development.

## Introduction

The novel coronavirus of 2019 (2019-nCoV), has affected nearly every corner of the world, affecting human health and causing enormous human loss (Seah and Agrawal, 2020). Originating in Hubei province, China, it causes severe respiratory problems in avians, as well as in mammals (Lorusso et al., 2020; Pal et al., 2020). This unprecedented epidemic has already been declared an international public health emergency by the World Health Organization (WHO). At the time of writing, the confirmed cases of this rapidly spreading coronavirus globally exceeded 12.6 million. Currently, there is no specific treatment for nCoV (Lu, 2020). Developing a new drug or vaccine is a major challenge; therefore, other antiviral drugs are fast-tracked for testing against nCoV (Li et al., 2020). Recent studies have shown that nCoV is mutating and evolving frequently (Zhang et al., 2020). Worldwide efforts were made to sequence the entire 2019-nCoV genome to study genetic variation and its evolutionary origin (Ceraolo and Giorgi, 2020).

Genome-level analysis will provide information on host responses and may improve vaccine development. 2019-nCoV is an RNA virus, has a higher rate of mutation that may have facilitated the widespread of this virus across different weather conditions (Pachetti et al., 2020). Information regarding the composition of amino acids and the codon usage pattern may help gain more insight into gene expression or other selection pressures. In this study, genetic variations of 2019-nCoV were analyzed based on available data. Here, we reported the emerging genetic variants that might have developed over time in humans since the virus became pandemic. To assess the genetic variation, one hundred and sixty four complete genome sequences of 2019-nCoV were retrieved from GISAID (Shu and McCauley, 2017). The genome sequence of Wuhan 2019-nCoV was used as a reference to identify the genetic diversity and mutations between the 2019-nCoV sequences from different countries. The hypervariable 2019-nCoV genomic region undergoing mutations were assessed for their tendency to alter the amino acid sequence in the respective polypeptide sequences.

## Materials and methods

SARS Cov-2 genomic sequences were retrieved from GISAID (Global Initiative on Sharing Avian Influenza Data) (Shu and McCauley, 2017). As of December 2019, GISAID became established as a database for the coronavirus. A total of 164 genomic sequences were collected for the present analysis, comprising 69 sequences from Indian states and 95 genomic sequences from different continents, including 62 countries. Here, we masked the initial 64 nucleotides and the sequences exceeding the position 29686 while performing multiple sequence alignment. ClustalW software was used for sequence alignment (Guo and Sun, 2000). The aligned sequences were used to create phylogenetic tree using MEGA X software (Kumar et al., 2018). The sequence data were analyzed using neighbor-joining method. To assess the robustness of individual nodes, the bootstrap method was employed to test the phylogeny with 1000 replicates (Soltis and Soltis, 2003). Sequence handling and data processing were carried out in Bioedit software (Hall et al., 2011). AutoDock 4.2 software was used to dock the protein structures with their potential ligands (Rizvi et al., 2013). Codon usage pattern of individual genomic sequence was determined based on RSCU analysis. INCA 2.1 software was used for codon usage as well as amino acid usage bias analysis (Supek and Vlahoviček, 2004).

## Results

### Phylogenetic Analysis

GSAID database was used to fetch 164 SARS CoV-2 genomic sequences which comprised of 69 genomic sequences from India and 95 from other countries of the World (**Figure. 1**) Genomic sequences from different Indian states have been phylogenetically analyzed to gain insight into the genetic variation prevalent in India (**Supplementary file 2: Figure 3**). Based on the phylogenetic tree we grouped the 2019-nCoV genome sequences of Indian states into 13 major groups (**Supplementary file 2: Figure 4**). These 13 representative groups (states) of India were then phylogenetically analyzed along with SARS CoV-2 genomes (95 coronavirus sequences) from different countries around the world. Based on phylogeny, 27 major groups which included 62 countries were created (**Supplementary file 2: Figure 5**).

**Figure 1:**
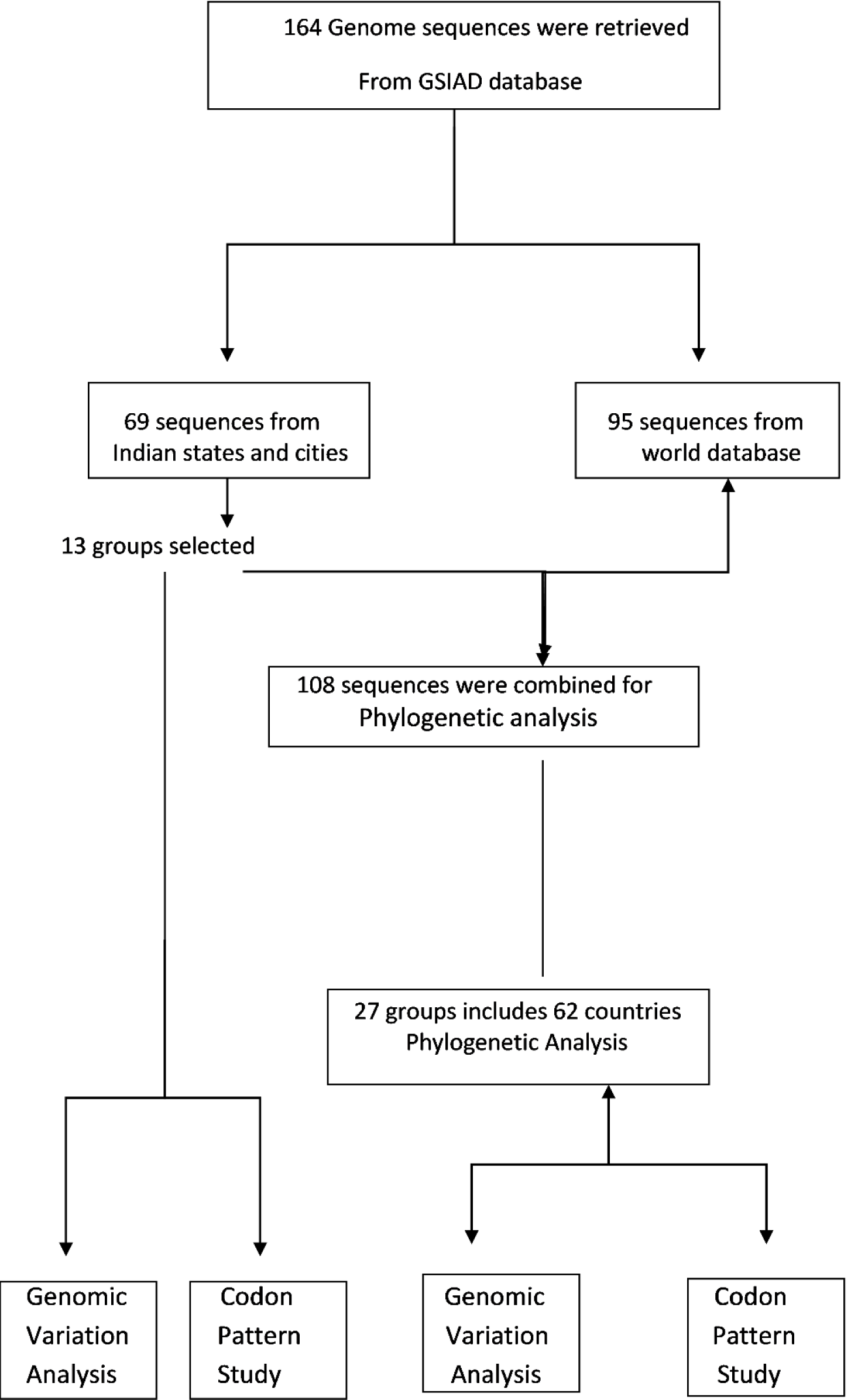
Flow chart representing the overall work done from retrieval to analysis of genomic sequences

### Analysis of genetic variation across the world

To investigate the genetic variation in the novel coronavirus, we compared the SARS-CoV-2 genomic sequence from 62 countries with the Wuhan SARS-CoV-2. Here we observed sequence conservation of 99.9 percent similar to previous findings (Lu et al., 2020). In comparison to the reference sequence (Wuhan strain), 83 single nucleotide polymorphism (SNP) was observed at various genomic locations in the coronavirus genome obtained from different countries. Among 83 SNP, maximum mutations were detected in orf1ab gene (37) followed by genomic region (26). The changes in the nucleotide has altered 32 amino acids whereas 25 mutations were recorded as synonymous mutations (**Table 1**).

**Table 1:** Single Nucleotide Polymorphism and amino acid changes identified in the entire Coronavirus genome (World Data)

| Genes | Total SNP | SNP led to change in Amino acid | Synonymous mutations |
| --- | --- | --- | --- |
| orf1ab | 37 | 16 | 21 |
| spike | 8 | 4 | 4 |
| Orf3a | 4 | 4 | 0 |
| Memberane | 3 | 3 | 0 |
| Orf8 | 3 | 3 | 0 |
| Nucleocapsid | 2 | 2 | 0 |
| Genomic | 26 |  |  |
| **Total** | **83** | **32** | **25** |

The description of genetic variation occurred in SARS CoV-2 genome with respect to each country (Selected for the present study) has been provided in **supplementary file 1: Sheet 1**. In addition, gene-wise mutational information for the selected countries have also been given **supplementary file 1: Sheet 2)**. For the present study, we focused on those regions which showed nucleotide variations in at least more than 6 genomes and are thus considered as hypervariable regions (**Table 2**). Hypervariable regions at 1068 and 11092 genomic position falls under the orf1ab gene (**Supplementary file 2: Figure 1**). At the amino acid level the mutation at 1068 (C→T mutation), and 11092 (G→T mutation) resulted in alteration of amino acid from threonine (ACT) to isoleucine (ATT) and asparagine (TTG) to phenylalanine (TTT), respectively. While at 3046 position, we observed a synonymous mutation. Another hypervariable region was located at genomic position 23412, this area encodes spike protein. Here we observed G instead of A, which leads to a shift in the encoded protein sequence, *i.e*. aspartate (GAT) to glycine (GGT). Variation at genomic position 25572 was localized in the gene orf3a. At this location, G→T mutation has resulted in glutamine (CAG) to histidine (CAT) substitution. Likewise, nucleotide (C→T) alteration was also observed in the nucleocapsid gene at the genomic position 28230 which resulted in a change in the amino acid (proline (CCC) was replaced by glutamate (CTC).

**Table 2:**
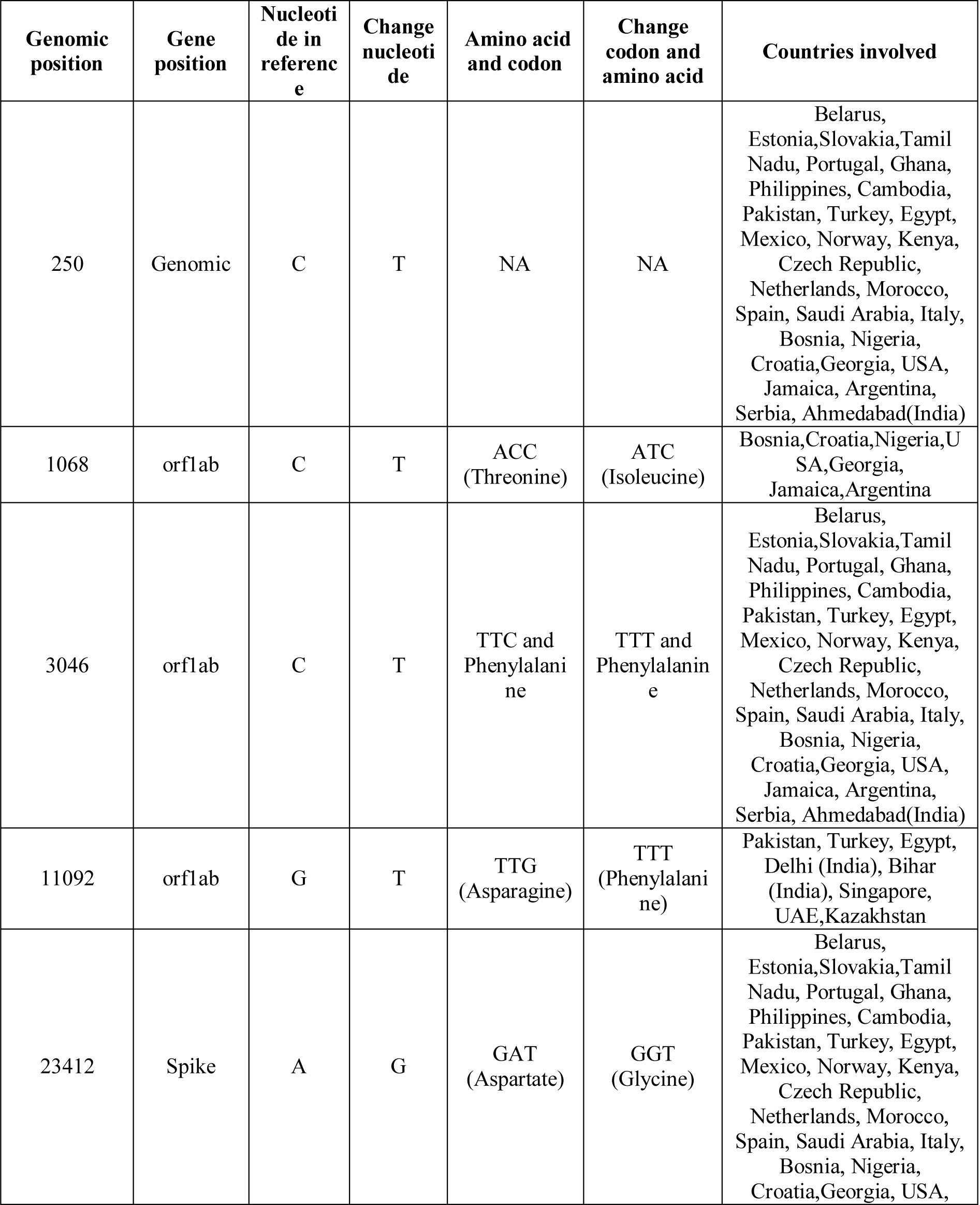
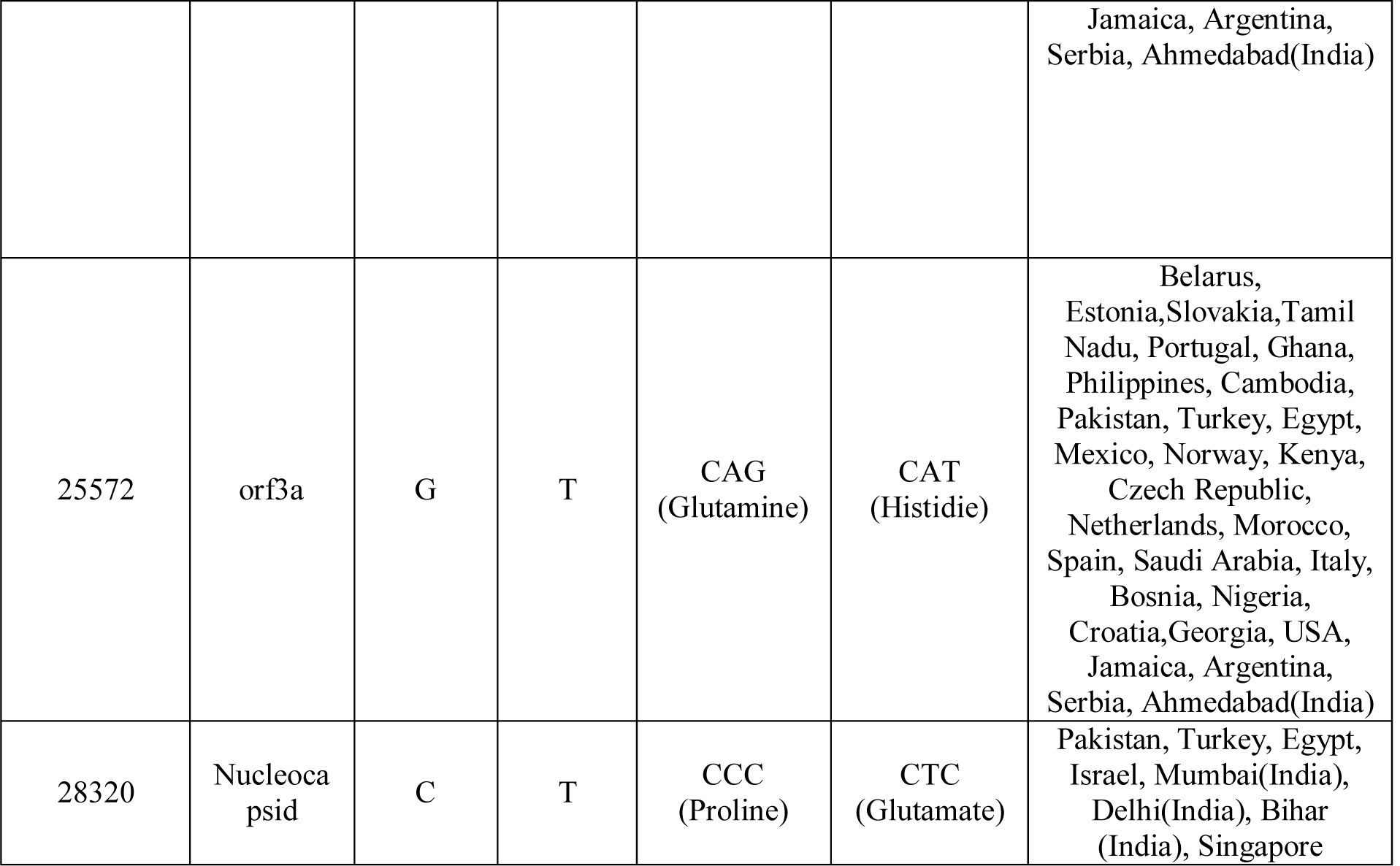
Hypervariable regions identified from entire SARS CoV-2 Genome (World Data)

### Analysis of genetic variation across Indian states

Genomic sequences of SARS CoV-2 obtained from different Indian states were also compared with the reference genome (Wuhan) to detect the genetic variation. A total of 47 SNPs were detected in 2019 nCoV-2 genomes obtained from Indian states. The implication of the nucleotide polymorphism has led to alterations in 21 amino acid while 16 mutations were recorded as synonymous (**Table 3**). State wise (selected states) description of genetic variations across the genome has been given in **supplementary file 1: Sheet 3**. Additionally, the number of mutations occurred in all the selected states with respect to their gene location has also been tabulated in **supplementary file 1: Sheet no 4**.

**Table 3:** Single Nucleotide Polymorphism and amino acid changes identified in the entire Coronavirus genome (World Data)

| Genes | Total SNP | SNP led to change in Amino acid | Synonymous mutations |
| --- | --- | --- | --- |
| orf1ab | 24 | 14 | 10 |
| spike | 8 | 4 | 4 |
| Orf3a | 2 | 1 | 1 |
| Memberane | 1 | 0 | 1 |
| Orf8 | 1 | 1 | 0 |
| Nucleocapsid | 1 | 1 | 0 |
| Genomic | 10 |  |  |
| **Total** | **47** | **21** | **16** |

For the present investigation, we have described the mutations that took place at hypervariable regions involving 3 or more than 3 genomes of Indian states (**Table 4**). We found four hypervariable genomic regions corresponding to orf1ab gene. These genomic positions are 3037, 6312, 11083, and 13730 Synonymous mutation was observed at 3037 (**supplementary file 2: Figure. 2**). The transition in amino acid from threonine (ACA) to lysine (AAA) was detected at position 6312 (non-synonymous mutation). At 1083 genomic position, there was a substitution of leucine (TTG) by phenylalanine (TTT) due to G →T mutation. At 13730, C→T mutation caused the alanine (GCT) to be changed to valine (GTT) in the polypeptide chain. In the genomic region of the spike gene at 23403 and 23929 locations, genetic variability was also identified. At 23403, non-synonymous mutation (GGA by GGT, *i.e*. AG mutation) was found forcing aspartate to alter in the final protein sequence by glycine. Likewise, amino acid modification from methionine (TAC) to tyrosine (TAT) occurred at 23929 position due to C→T mutation. Likewise, C→T mutation in the nucleocapsid gene (at 28311 genomic position) results in a change in amino acid, glutamate (CTC) replaced proline (CCC).

**Table 4:**
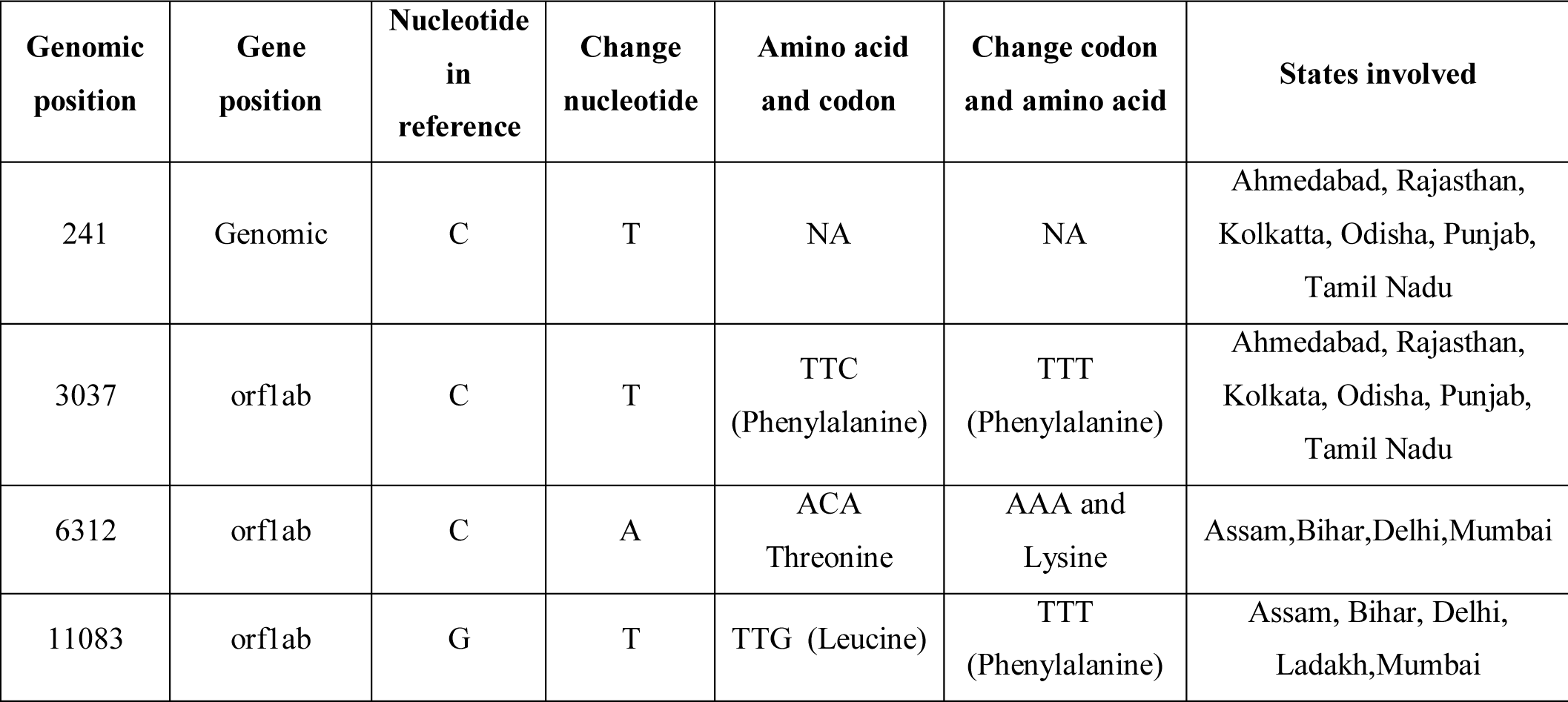
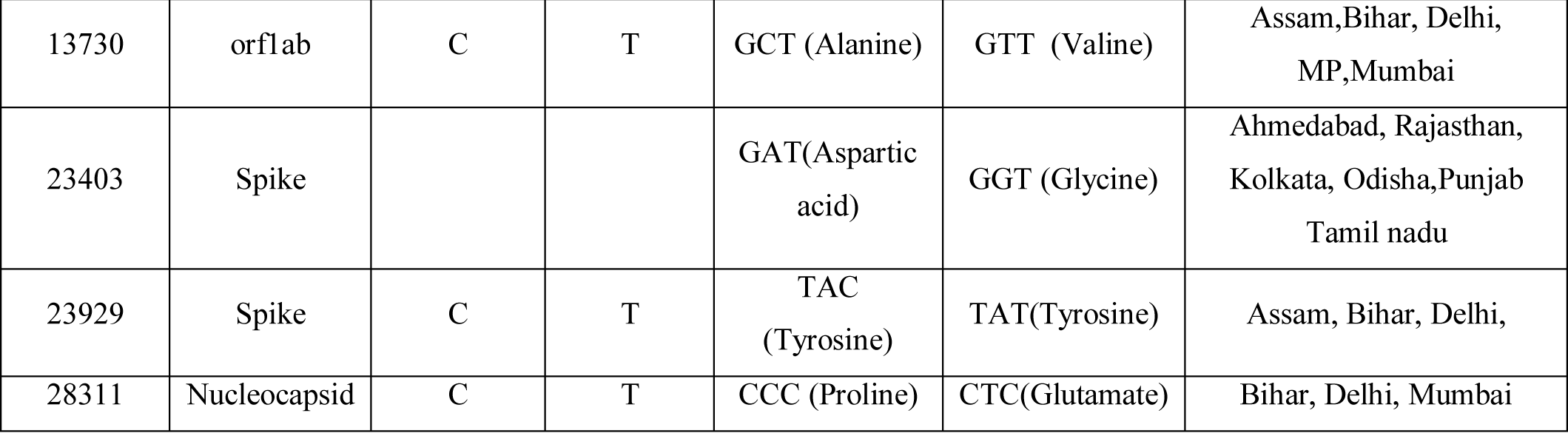
Hypervariable regions identified from entire SARS CoV-2 Genome (Indian States)

### Worldwide: Codon usage and amino acid usage bias

Genome of coronavirus sequenced from 62 countries (27 groups based on phylogeny) were analyzed for codon usage trend and amino acid composition. Based on the similarity in the pattern of codon usage and amino acid composition among 27 groups, these were further assembled into 3 groups A, B, and C. Group 1 to 4 formed Group A, Group 5-7 clustered into Group B and Group 8 onwards constituted group C. We observed that coronavirus genome in all the 3 groups showed a similar pattern in codon usage with respect to 7 amino acids Cys, Glu, Phe, Ileu, Lys, Leu, and Gln that preferred to use U/A ending codons. Differences in the Codon usage pattern for 11 amino acids were observed among the 3 groups. SARS CoV-2 genome of group A showed a tendency to use U-ending codons for 10/11 amino acids except for arginine (AGG). Genomes from group C also preferred to use U-ending codons for 8/11 amino acids except proline, arginine, and threonine that tends to use A-ending codons. Whereas the coronavirus genomes of group B displayed a distinct pattern of codon usage by choosing A and C ending codons. Moreover, uniqueness in the use of G-ending codon for valine was also detected in the genomes of group B. The data has been provided in (**supplementary file1: Sheet no 7**). Amino acid composition analysis revealed that leucine was found to be predominant among all three groups however, differences were observed in the usage frequency. Group B showed maximum frequency (0.16) as compared to Group A (0.12) and C (0.08) (**supplementary file 1: Sheet no 8**).

### Indian States: Codon usage and amino acid usage bias

Coronavirus genomes from 13 selected Indian states (Based on phylogeny) have been studied for their variability in codon usage. Diversity in the usage of synonymous codons were observed in the genomes sequenced from different states. We noticed that nine states (Ahmedabad, Assam, Bihar, Rajasthan, Delhi, Ladakh, Mumbai, Odisha and Punjab) exhibited identical trends in codon usage. Madhya Pradesh (MP) and Tamil Nadu (TN) showed a similar trend, while Kerala and Kolkata showed a distinct pattern compared to other states. We also observed that SARS CoV-2 genomes from all states prefers U-ending codons to encode majority of the amino acids except Leu, Lys, Gln, Arg, Ser, and Thr, which preferred to be coded by A-ending codons. Genomes obtained from different states demonstrated bias in codon selection to encode three amino acids, namely serine, tyrosine, and valine. Genomes sequenced from Kerala, Kolkata, MP, TN prefer to use the UCA codon to encode serine, while genomes from other states tend to use AGU codon. Interestingly, genomes sequenced from Kerala showed heterogeneity in tyrosine (UAC) and valine (GUG) codon selection as opposed to strains isolated from other states (**supplementary file 1: Sheet 5**).

Compositional analysis of amino acids showed the dominance of leucine in all the 2019-nCoV genomes analysed. However, variability in the frequency of use of leucine was observed among the genomes retrieved from different states. Kerala showed the highest frequency (0.16) followed by Kolkata, MP, and TN (0.12) and then the rest of the states (0.08). It has been observed that 6-fold and 4-fold degenerative amino acids are more common compared with 2-fold amino acids. Glutamate, aspartate, and proline were among the lowest frequency amino acids in all the coronavirus genomes (**supplementary file 1: Sheet 6**).

## Discussion

Rapid evolution and spread of the corona virus across the world intrigued us to understand the genetic variation in the SARS CoV-2 genome. Our findings also revealed the existence of high mutation rates in 2019-nCoV genomes sequenced from different parts of the world. Moreover, phylogenetic analysis further confirmed the migratory context of 2019-nCoV and its spread. The hypervariable regions identified in the SARS CoV-2 genes (orf1ab, S and nucleocapsid N gene) also supports the constant evolution of this virus across the world. In orf1ab gene, we found two hypervariable amino acid changing region (1068 and 11092 genomic locations) worldwide and three hypervariable regions (6312, 11083, and 13730 genomic locations) in Indian states when compared with the Wuhan strain. Single nucleotide polymorphisms were observed highest in Orf1ab. The mutations reported in the Orf1ab gene will be of interest to investigate the clinical function of this protein in Covid-19. It has been reported earlier that ORFs and ACE2 genes play a key role in the establishment of the pathogenesis of coronavirus (Koyama et al., 2020).

In addition Nucleocapsid N gene also showed a hypervariable region where proline was replaced by glutamate due to mutation. The N protein plays a key role in virion assembly (McBride et al., 2014). Therefore, this hypervariable region might have influenced its assembly efficiency and thus pathogenicity. Similarly, we observed a hypervariable region in S protein where (due to mutation) aspartate was replaced by glycine. In addition, across the Indian states, we found another non-synonymous mutation in S protein where methionine was replaced by tyrosine. Non-synonymous mutation affects the overall conformation of proteins. S protein plays a crucial role in viral infection and pathogenesis (Phan, 2020). Additionally, we observed altered binding confirmation when we docked the mutated S protein with ramdesivir and NAG compared to the Wuhan S protein (**supplementary file 2: Figures 6 & 7**). We may, therefore, believe that this altered amino acid may have made a major contribution to the pathogenesis and transmission of 2019-nCoV.

Since the COVID-19 outbreak in Dec 2019, large number of studies has reported the presence of SNPs in different genomic locations. Here we observed 37 SNPs at different genomic locations encoding ORF1ab gene. Genome wide comparison showed that Denmark, England and Ahmedabad (India) has the maximum mutation in ORF1ab gene compared to all the countries analyzed. Banerjee and colleagues reported 14 SNPs in the ORF1ab gene (Banerjee et al., 2020). Similarly, Ceraolo and Giorgi 2020 revealed the presence of one hypervariable region in ORF1ab (Ceraolo and Giorgi, 2020). Previous research on S protein isolated from India showed the presence of single mutation when compared with the Wuhan strain (Saha et al., 2020). Tunisia, Iran showed 2 noteworthy mutation in the genomic locations corresponding to S protein. Similar findings were observed for the following Indian states namely Ahmedabad, Bihar, Rajasthan, Kolkata and Punjab. In addition to the mutation in the coding region we also detected 26 mutations in different genomic locations of SARS-CoV-2.

Furthermore, genome-wide codon usage comparison between Indian sates revealed the bias in codon selection to encode three amino acids, namely serine, tyrosine, and valine. Likewise, we observed eleven amino acids namely, Ala, Arg, Asn, Asp, Gly, His, Pro, Ser, Thr, Tyr, and Val showed variation in codon selection between world genomes divided into three groups. These findings, which reflect substantial differences in the use of codon between the 2019-nCoV quasispecies, will be useful for assessing the adaptation, evolution of a host-virus and are therefore of interest to vaccine design strategies.

In conclusion, our analysis confirms high variability among 2019-nCoV sequenced quasispecies, highlighting hypervariable positions within three key protein-coding regions. Such variability in proteins might have affected the patient’s clinical outcomes because the viral genome that infects them is slightly different. Nevertheless, there is a chance that these mutational variants might have modulated the disease’s spread. This is the first report on the genetic variation present in different states of India in comparison with other countries. Our results provide positive light on the prospect of developing 2019-nCoV therapy for patients from different locations.

## Authors contributions

All the authors carried out the analysis, RS and PP conceived the idea. SG prepared display items. SG and RS wrote the research article. PP edited the final manuscript. All the authors reviewed the manuscript before submission.

## Conflict of interest

None declared

## Acknowledgements

Authors are thankful to Negenome Bio Solutions group for their support. Authors are also grateful to the IIIM Jammu for providing the necessary research facilities to carry out this work.

## Notes

### Competing Interest Statement

The authors have declared no competing interest.

