## Supplementay file 2 for "Analysis of single nucleotide polymorphisms between 2019-nCoV genomes and its impact on codon usage"

**Manuscript Details:**

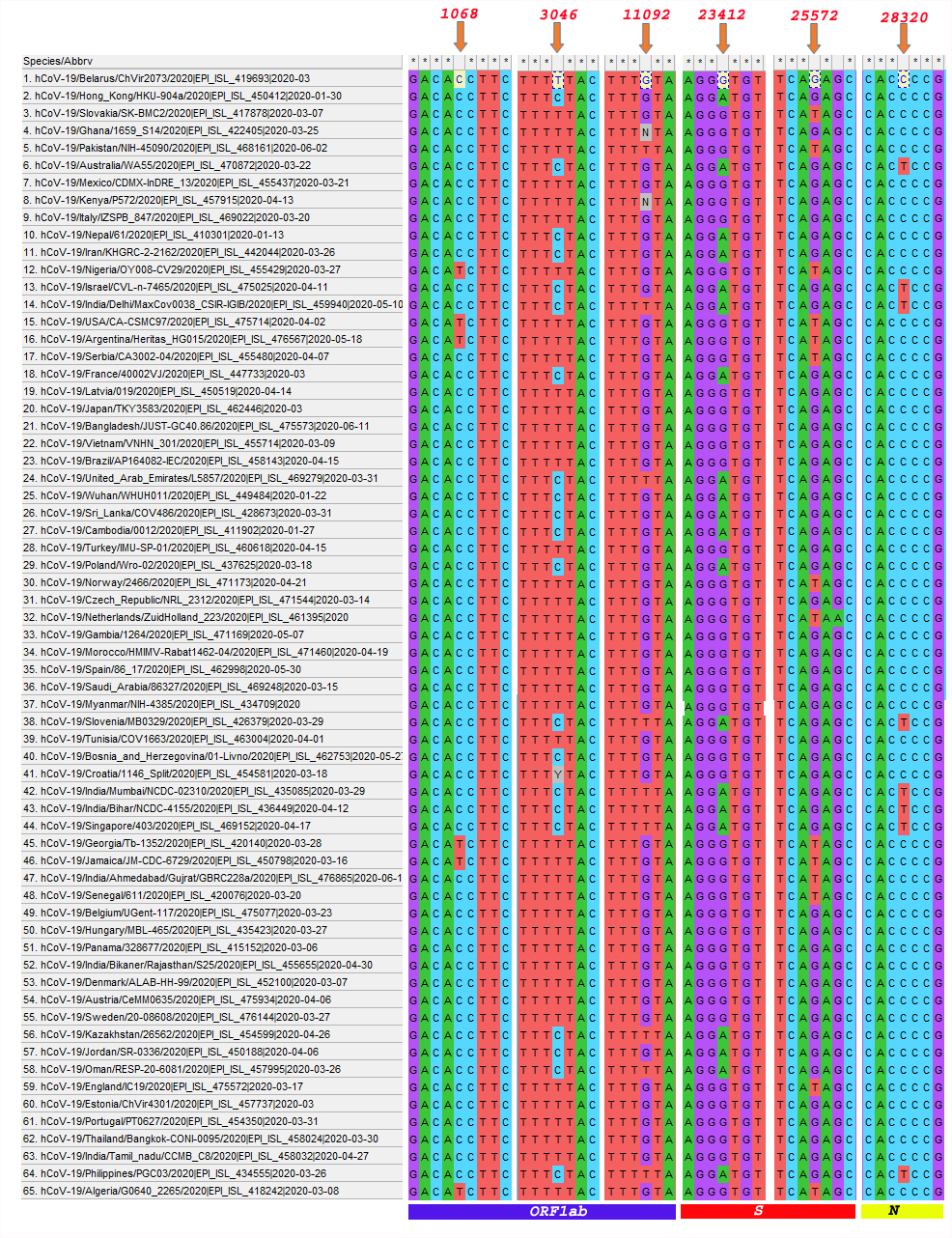

**Fig 1: Multiple sequence alignment 65 world SARS CoV-2 genomes (including 5 Indian States) representing the hypervariable regions across the entire genome. (Mutations at 1068bp, 3046bp and 11092bp genomic positions are localized in gene orf1ab), (Mutations at 23412 bp and 25572bp are localized in spike gene), (Mutations at 28320 bp are localized in Nucleocapsid (N) gene)**

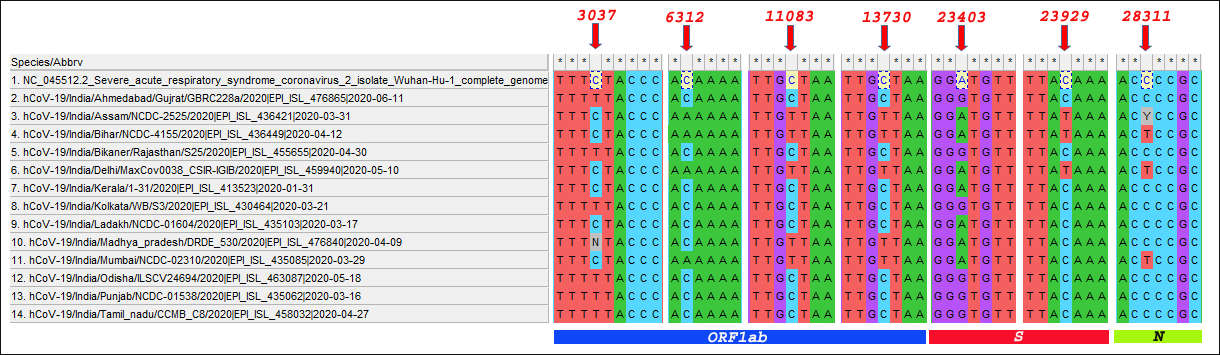

**Fig 2: Multiple sequence alignment of SARS CoV-2 genomes from selected 13 Indian representing the hypervariable regions across the entire genome (Mutations at 3037bp, bp and 6312bp, 11083bp, 13730bp genomic positions are localized in gene orf1ab), (Mutations at 23403 bp and 23929 bp are localized in spike gene), (Mutation at 28311 bp are localized in Nucleocapsid (N) gene)**

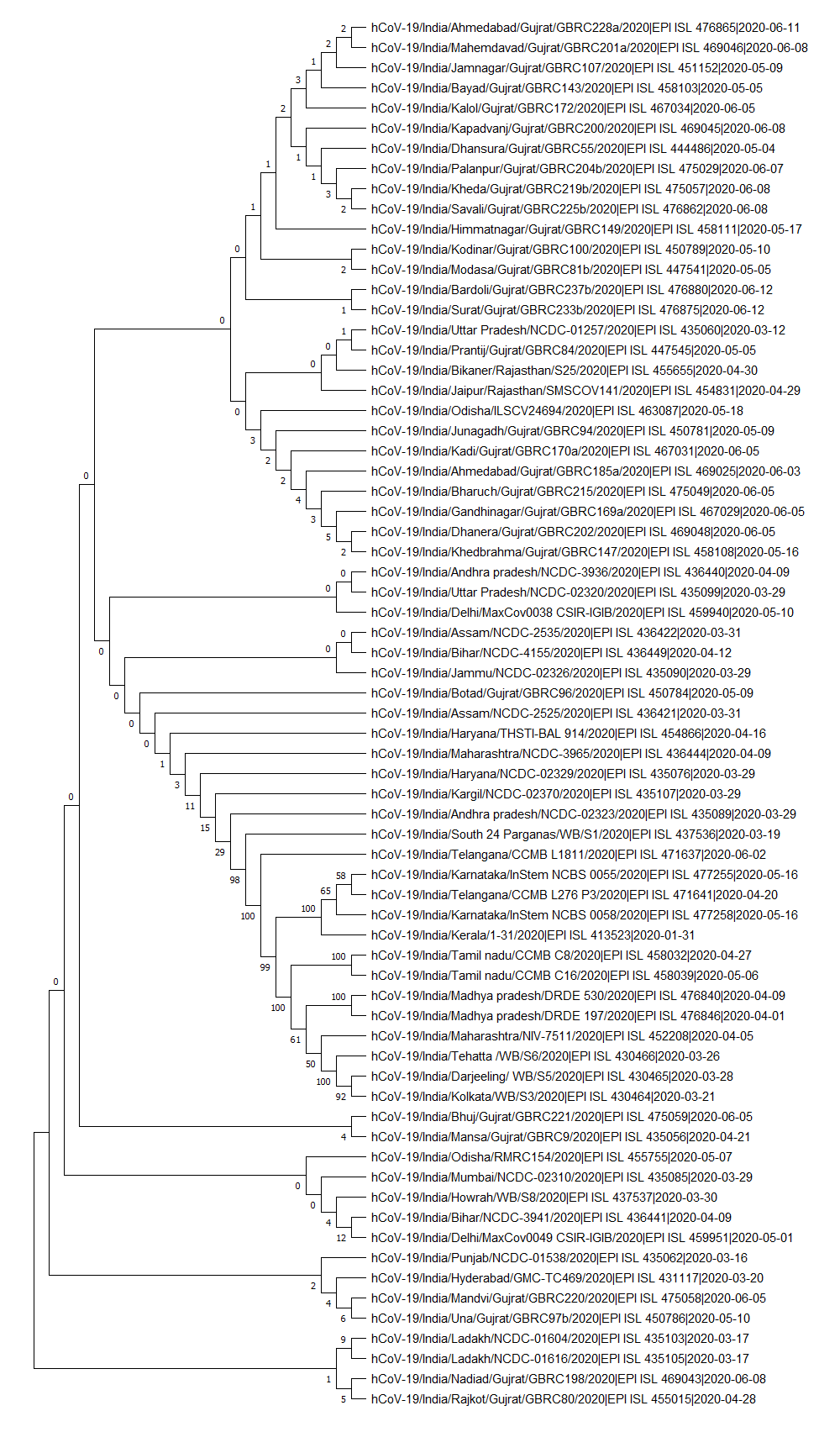

Fig 3: Phylogenetic tree of 69 SARS CoV-2 genomes representing different cities/ states of India

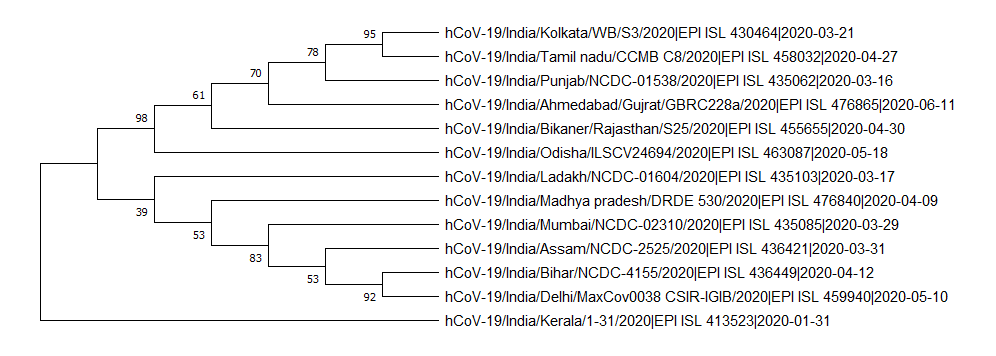

Fig 4 : Phylogenetic tree of SARS CoV-2 genome sampled from 13 selected Indian states

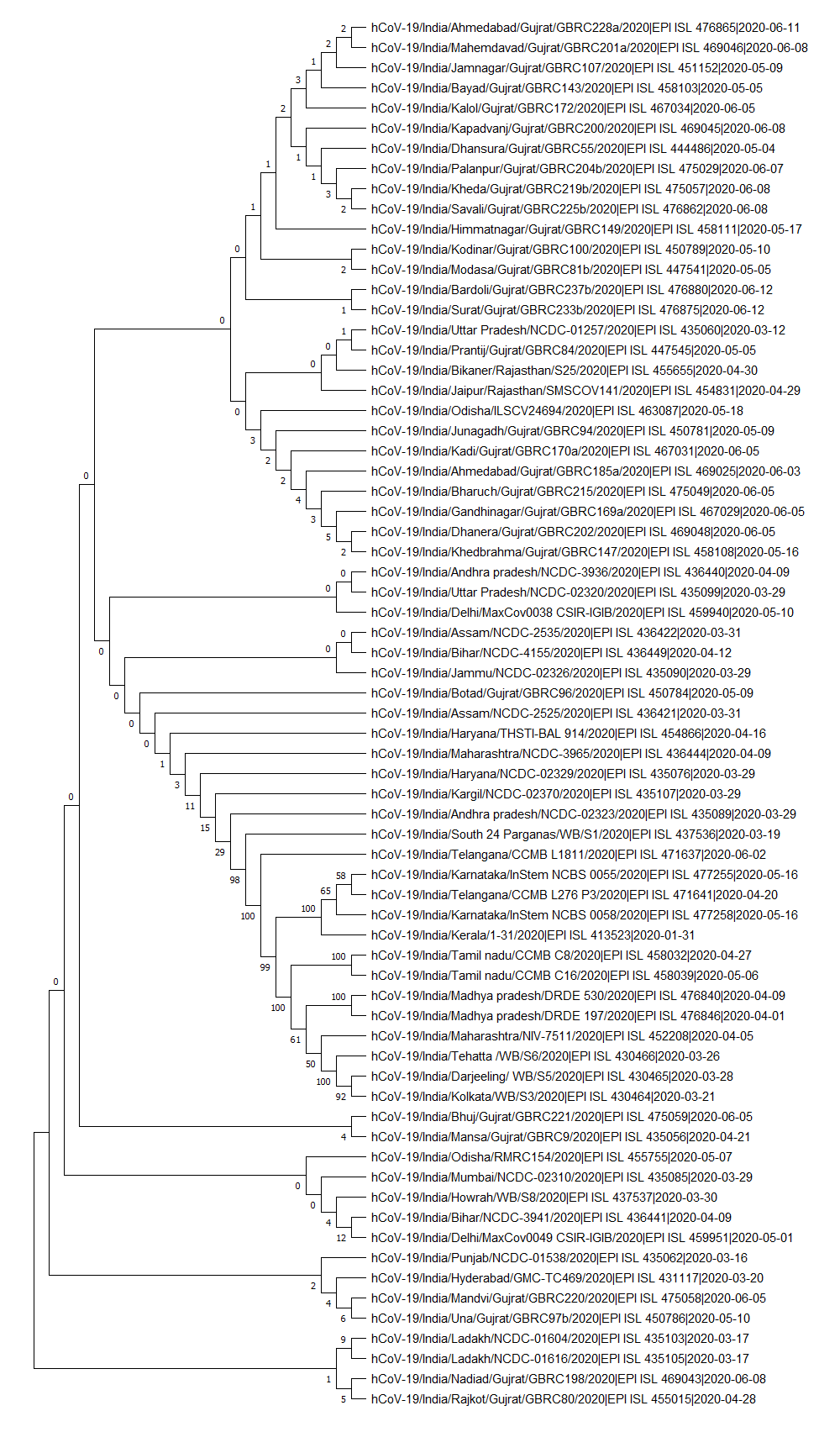

Fig 5: Phylogenetic tree constructed using 108 (95 global and 13 Indian states) SARS CoV-2 genomes

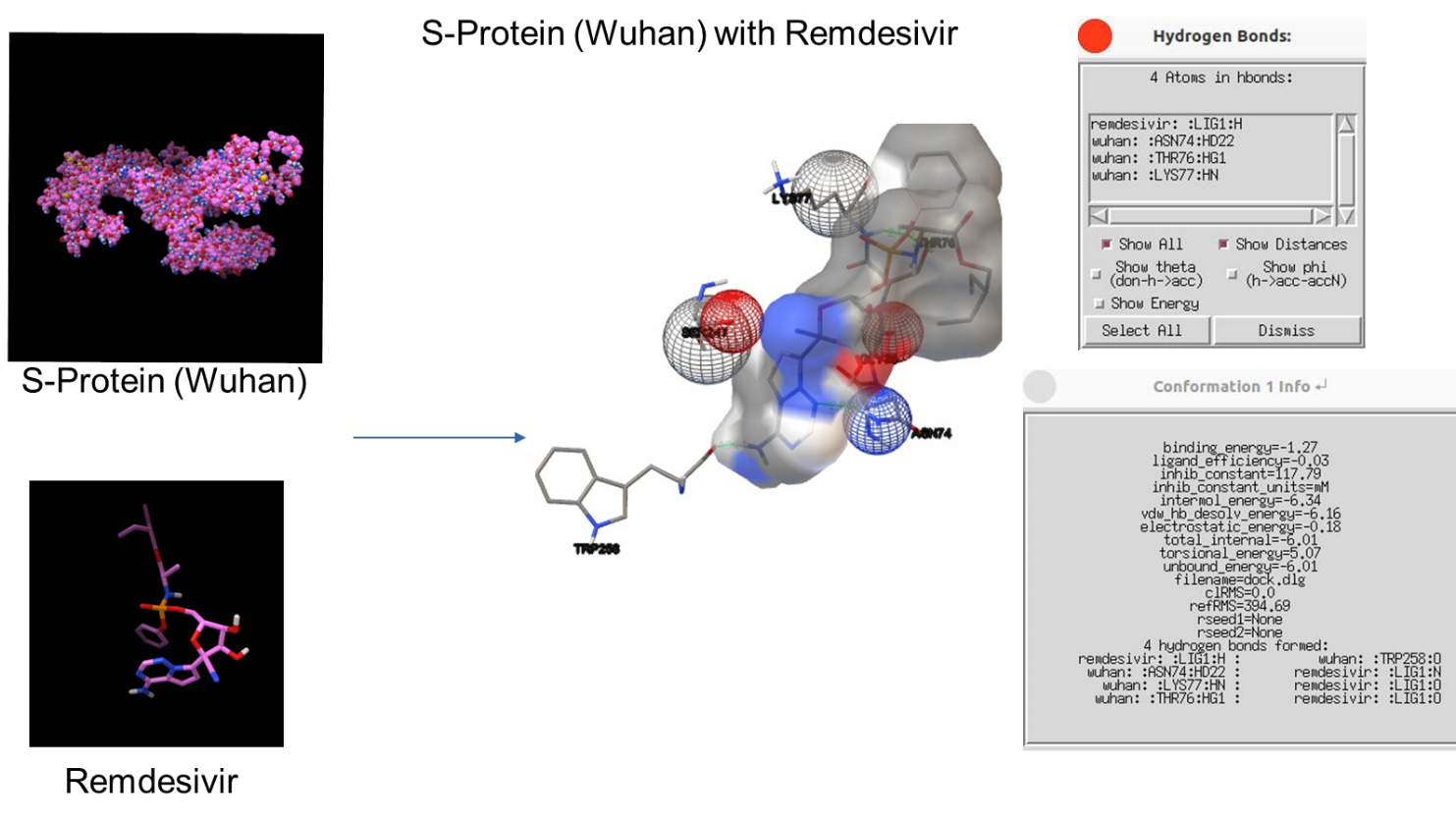

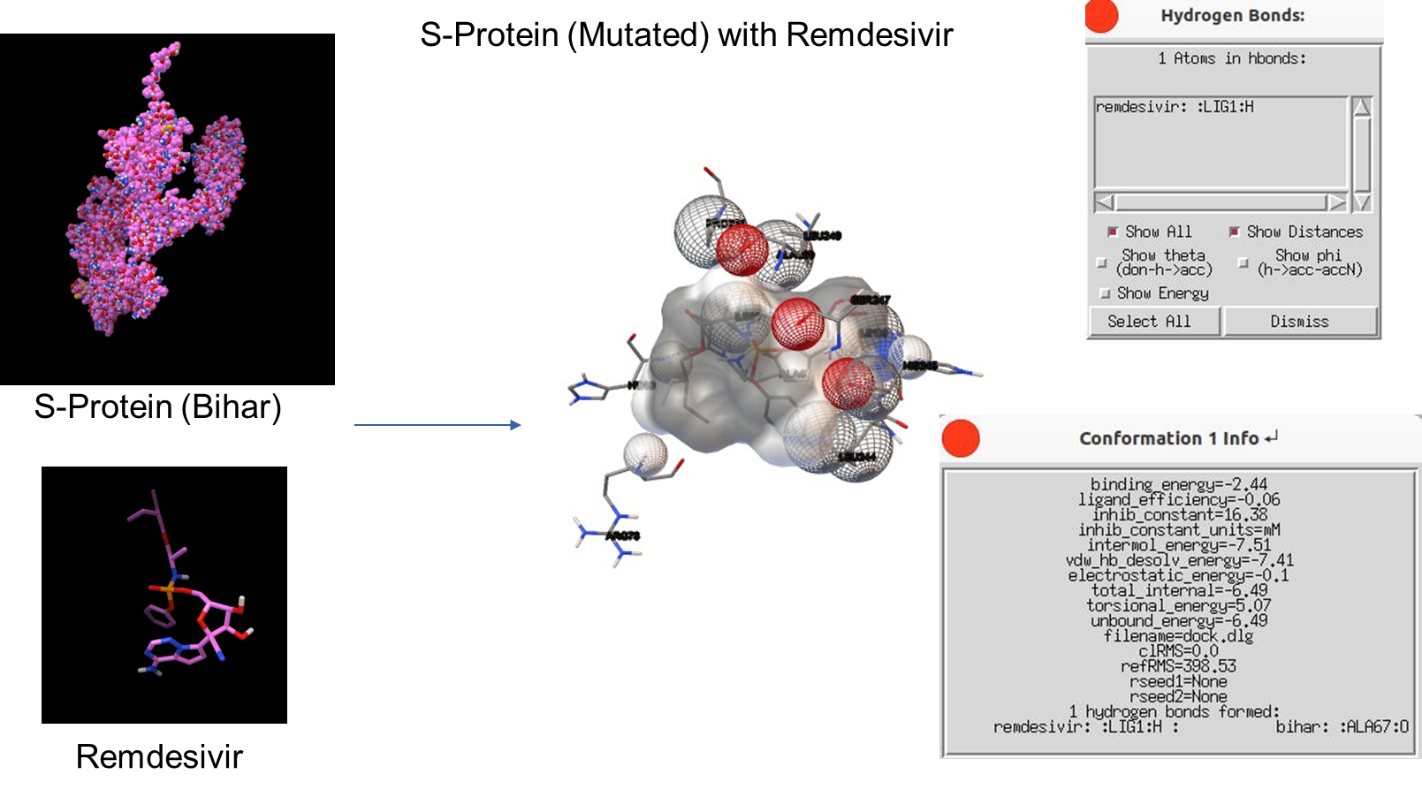

**Fig 6:** Docked structure of S protein with Remdesivir

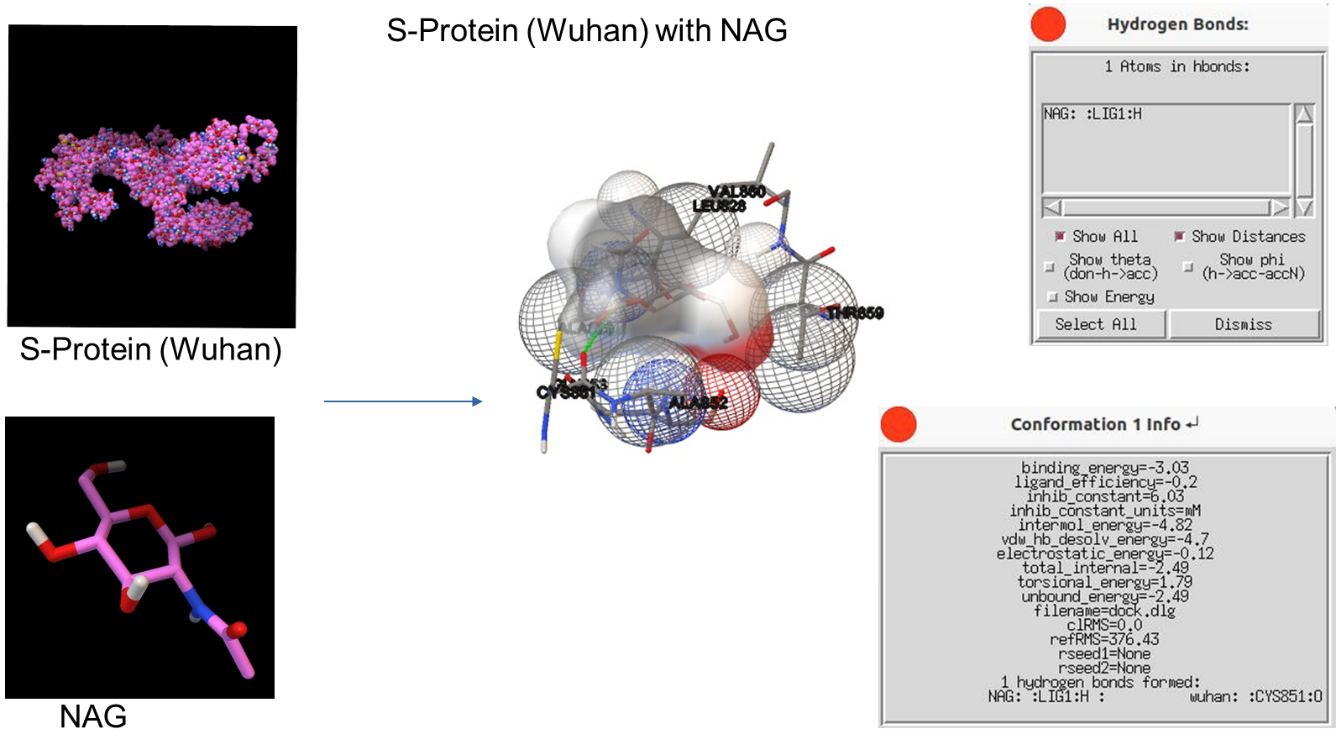

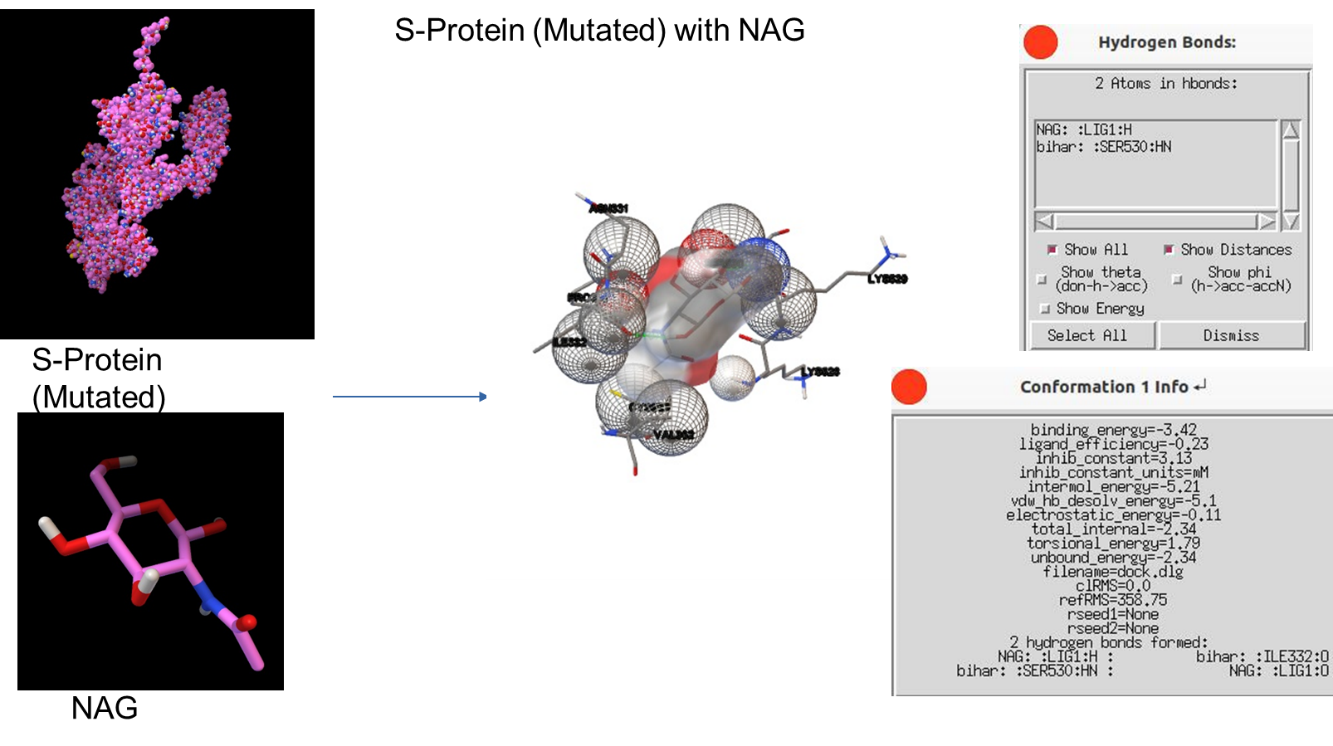

**Fig 7:** Docked structure of S protein with NAG
